# Chromosome-level reference genome assembly for the mountain hare (*Lepus timidus*)

**DOI:** 10.1101/2024.06.10.598177

**Authors:** Zsófia Fekete, Dominic E. Absolon, Craig Michell, Jonathan M. D. Wood, Steffi Goffart, Jaakko L. O. Pohjoismäki

## Abstract

We present here a high-quality genome assembly of a male mountain hare (*Lepus timidus* Linnaeus), from Ilomantsi, Eastern Finland, utilizing an isolated fibroblast cell line as the source for high quality DNA and RNA. Following the previously published brown hare reference genome assembly, the mountain hare is the second Finnish pilot species for the European Reference Genome Atlas (ERGA) initiative, a collaborative effort to generate reference genomes for European biodiversity.

The genome was assembled using 21× PacBio HiFi sequencing data and scaffolded using the Hi-C chromosome structure capture approach. After manual curation, the primary assembly length was 2,695,305,354 bp with N50 125,755,317 bp. The largest scaffold was 181 Mbp and the scaffold N50 127 Mbp, contributing to a primary assembly consisting of 85 scaffolds and an alternate assembly with 109 scaffolds. The scaffolds include 23 autosomes, numbered according to their size, as well as X and Y chromosomes, matching the known karyotype. Telomeric regions were present on at least one end of 19 of the chromosomes. The genome has a high degree of completeness based on the BUSCO score (mammalia_odb10 database), Complete: 95.1 % [Single copy: 92.3 %, Duplicated: 2.7 %], Fragmented 0.8 %, and Missing 4.1 %. The mitochondrial genome of the cell line was sequenced and assembled separately. The assembly meets the Earth BioGenome Project criteria for a reference-standard genome assembly.

Compared to the previous pseudo-reference genome assembly of *L. timidus* ssp. *hibernicus* Bell, assembled using the rabbit genome, this new reference genome represents the nominate subspecies and the species-specific chromosomal conformation. The published genome assembly will provide a solid foundation for future genomic research on Lagomorpha, including the insights into the genomic basis of adaptations to snowy and cold environments. Furthermore, it opens opportunities for experimental analysis of mountain hare gene functions.

## Introduction

The mountain hare (*Lepus timidus* Linnaeus, 1758), including several subspecies, is the most widespread hare species in the world, with its distribution ranging from Ireland to Japan (Angerbjörn & Schai-Braun, 2022). The species is adapted to the cold and snowy conditions of northern Eurasia by having wide snowshoe feet and white winter pelage. The ongoing climate change-induced shortening of the snow-covered season is detrimental for the mountain hare, especially due to camouflage mismatch (Zimova et al., 2016; Zimova et al., 2020), resulting in range contraction to the North or higher altitudes in mountains, while benefitting the expansion of its more southern relative, the brown hare (*L. europaeus*) (C.-G. Thulin, 2003; Jansson & Pehrson, 2007; Reid, 2011; Levänen, Kunnasranta, et al., 2018; Rehnus et al., 2018).

The mountain hare has shown resilience during the past warm periods of the Holocene (Smith et al., 2017), diverging also to more temperate climate-adapted subspecies (Giska et al., 2022). Notable examples include the Irish hare (*L. t. hibernicus* Bell, 1837) and the heath hare (*L. t. sylvaticus* Nilsson, 1831), which are likely relatively old evolutionary lineages that are genetically distinct from the widespread nominate subspecies (Giska et al., 2022; Michell et al., 2022). In addition, post-glacial contact with other congeneric hare species has resulted in recurrent hybridization and genetic introgression (Melo-Ferreira et al., 2009; Levänen, Thulin, et al., 2018), which appears to have adaptive significance (Giska et al., 2019; Pohjoismäki et al., 2021; Giska et al., 2022).

Besides the fact that geographically restricted, relic-like subspecies are threatened (Caravaggi et al., 2017; C. G. Thulin et al., 2021) and thus interesting in the context of conservation genetics, hares provide ample opportunities to study local adaptations (Giska et al., 2022; Michell et al., 2022), genomic introgression (Levänen, Thulin, et al., 2018; Giska et al., 2019; Pohjoismäki et al., 2021) and genetic constituents of species boundaries (Gaertner et al., 2023). Such studies are greatly facilitated by the availability of reference-level genome assemblies (Michell et al., 2024).

Relatively recent advances in sequencing technologies, so-called third generation sequencing or high-throughput sequencing of long molecules, has provided affordable means to produce highly continuous genome assemblies. Combined with techniques for sequence contact maps for chromosomal scaffolding, these methods enable reference-level genome assemblies for almost any species (Lawniczak et al., 2022), a level previously achieved only by a few model organisms. This in turn has resulted in emergence of several sequencing initiatives to produce reference genome assemblies for various taxa, including the European Reference Genome Atlas (ERGA) and the Darwin Tree of Life (DToL) (Blaxter et al., 2022). In the presented work, we use these methods to provide a reference-level sequence assembly for the mountain hare. Together with our previously released brown hare genome assembly (Michell et al., 2024), the mountain hare genome represents another Finnish contribution to the ERGA pilot project, aimed to demonstrate the feasibility of a continent-wide collaboration to sequence and assemble reference genomes for local species. Through resource and knowledge sharing, this collaborative approach not only accelerates the creation of reference-grade genomes but also enhances biological discovery, benefiting the broader scientific community (Mc Cartney et al., 2024).

Since long-molecule sequencing relies on intact, high-molecular-weight DNA, requiring fresh or flash-frozen tissue samples, we used a living fibroblast cell line derived from a male mountain hare specimen from Ilomantsi, Eastern Finland (Gaertner et al., 2023), as the DNA source. The sex of the specimen was important to cover the Y-chromosome. The sampling locality was chosen outside of the brown hare range to minimize the probability for hybridization (Levänen, Kunnasranta, et al., 2018; Levänen, Thulin, et al., 2018). Furthermore, the sampled population represents a continuum across the vast eastern Taiga zone, making our sample very representative geographically for the nominate mountain hare subspecies. It is worth noting that Linné did not formally designate a type specimen for the mountain hare, and the type locality has been later decided to be Uppsala in Sweden (Angerbjörn & Schai-Braun, 2022). The Swedish hares belong to the same larger, geographically connected Fennoscandian population as our specimen (Levänen, Thulin, et al., 2018; Michell et al., 2022). This is important because sampling from or near the type locality is considered desirable for the representativeness of the reference genome assembly (Lawniczak et al., 2022).

Following the same approach as with our brown hare reference genome assembly (Michell et al., 2024), we assembled a highly complete genome of the nominate mountain hare subspecies (*Lepus timidus timidus*) using 21× genome coverage of PacBio HiFi read data. The 2.69 Gbp assembly was further scaffolded with Hi-C sequencing data to chromosome-level, including all of the expected 23 autosomes, and X and Y sex chromosomes (2n = 48) (Gustavsson, 1972). Telomeric regions were present on at least one end in 19 of the assembled chromosomes, further demonstrating the high continuity of the assembly.

While there is an existing genome assembly for the mountain hare (GCA_009760805), it represents the Irish subspecies *L. t. hibernicus* (hereafter the “Irish hare”) and is based on a female specimen, thus lacking the Y-chromosome. Also, the Irish hare genome assembly can be considered to be a “pseudoreference”, as it has been scaffolded using the rabbit (*Oryctolagus cuniculus* L.) reference genome assembly (Marques et al., 2020), and retains the rabbit 2n = 44 karyotype (Korstanje et al., 1999). Compared to the previous, our genome assembly represents a significant improvement both in continuity and scaffolding, providing a robust reference for future studies on adaptation, hybridization, and conservation of the mountain hare. Additionally, it can offer insights into the evolutionary history of hares, genetic basis of physiological adaptations, as well as expand the molecular toolkit for experimental investigation of genes and their functions.

## Methods

### Sampling and confirming of the species identity

A young male mountain hare was sampled during a routine hunt by the last author in October 2018 at Kelovaara, Ilomantsi, Finland (63.0609 N, 30.6248 E). The sampling had minimal impact on the local mountain hare population and no impact on the habitat. As mountain hares are legal game animals in Finland and the hunting followed the regional hunting seasons and legislation (Metsästyslaki [Hunting law] 1993/615/5§), the sampling adheres to the ARRIVE guidelines and no ethical assessment for the sampling was required. Sampling also did not involve activities governed by the Convention on International Trade in Endangered Species of Wild Fauna and Flora (CITES) or other export of specimens, as defined by the Convention on Biological Diversity (CBD).

The species identity was confirmed at site based on the morphological features distinguishing the mountain hare from the brown hare, the only other hare species in northern Europe. The collection location is remote wilderness, dominated by boreal Taiga Forest (Figure 1A) and devoid of brown hares, which in Finland are dependent on man-made environments (Levänen, Kunnasranta, et al., 2018). Further analysis of the coding part of the genome and mitochondrial DNA haplotyping showed no ancestral admixture with brown hares (Gaertner et al., 2023).

**Figure 1.**
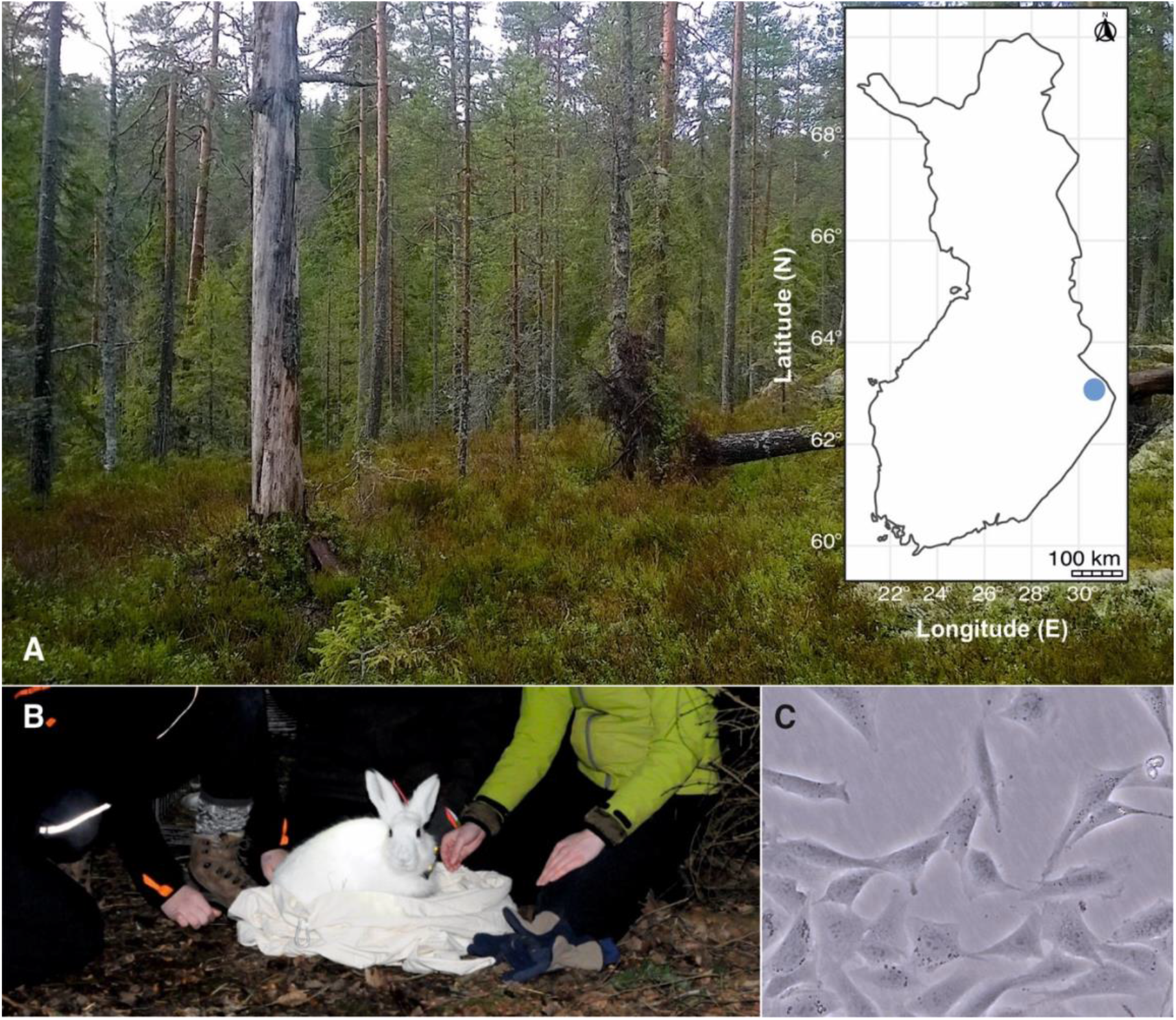
A) The geographic location in Finland and a view of the sampling biotope in Ilomantsi. Photo by Jaakko Pohjoismäki. B) A male mountain hare caught in the same region. Although the cell line used in this study originates from a killed specimen, fibroblasts can be isolated from sources such as ear clippings taken during tagging of animals for behavioral studies. Photo by Mervi Kunnasranta. C) Phase contrast microscopic image of the LT1 cell line used for the genome sequencing and assembly. The cells exhibit a typical fibroblast morphology. Photo by Steffi Goffart.

### Generation and vouchering of the cell line

We generated the fibroblast cell line (LT1) from a skin biopsy obtained from the specimen as described earlier (Gaertner et al., 2023). The primary cells were subsequently immortalized through SV40 large T-antigen transformation to enable their long-term maintenance for experimental use. The resulting cell line has been deposited under voucher number ZFMK-TIS-69753 in the biobank of the Stiftung Leibniz-Institut zur Analyse des Biodiversitätswandels, Zoological Research Museum Alexander König (ZFMK), Bonn, Germany, ensuring accessibility for future research. For genome sequencing and assembly, DNA was extracted from cells at passage number 10, as counted from the initial isolation of the primary cells. Typically, cell lines with under 20 passages are considered to have a low passage number (ATCC, 2024) and thus also having low risk of genomic changes associated with prolonged cell culture.

### High molecular weight DNA extraction and PacBio HiFi sequencing

High molecular weight DNA was purified using the classical phenol:chloroform:IAA extraction method (Sambrook & Russell, 2006). Briefly, the cells were grown to confluency on a 10 cm cell culture dish and detached using trypsin, followed by centrifugation and two washes with PBS. The cell pellet was resuspended in 2 ml of DNA extraction buffer (25 mM EDTA pH 8.0, 75 mM NaCl, 1 SDS and RNAseA), followed by Proteinase K digestion and phenol extraction, as detailed in the protocol (Sambrook & Russell, 2006). As done previously (Michell et al., 2024), the sequencing library preparations and sequencing on the PacBio Sequel II was performed in the DNA Sequencing and Genomics Laboratory, Institute of Biotechnology, University of Helsinki. Briefly, the DNA was quantified using Qubit and the fragment length was assessed with the Fragment Analyzer (HS large fragment kit) and then sheared to a length of 24 kb using the Megaruptor 3 (Diagenode) (45/200/30). Following this, the buffer was replaced with PacBio’s Elution buffer using AMPure beads. The DNA underwent repair and A-tailing, adapter ligation, nuclease treatment, and cleanup with SMRTbell cleanup beads, utilizing the SMRTbell prep kit 3.0. Fragments larger than 10,000 bp were purified with the BluePippin (Sage Science) (0.75% DF Marker S1 high pass 6-10kb v3), and the DNA was further purified using PacBio AMPure beads. It was then treated with DNA damage repair mix (PacBio) at 37°C for 60 minutes, followed by another purification with AMPure cleanup beads and elution into 11 μl of PacBio’s Elution buffer. Libraries were sequenced on two SMRT flow cells of the PacBio Sequel II at a concentration of 90 pM according to PacBio’s instructions, as provided by the SMRTlink software.

### Mitochondrial DNA (mtDNA) sequencing

The mitochondrial genome of this hare cell line has been sequenced for the purposes of another study by PCR amplification of approximately 2 kb overlapping mtDNA fragments (Tapanainen et al., 2024) (GenBank accession # OR915850).

### Hi-C library preparation

As with the brown hare genome assembly (Michell et al., 2024), Hi-C sequencing libraries were prepared following the protocol of (Belaghzal et al., 2017) with the following changes: 1.) The Hi-C protocol was performed in triplicates to produce a diverse sequencing library; 2.) NEBNext Ultra II FS DNA module was used for size fractionation; 3.) the NEBNext Ultra II library preparation kit for Illumina was utilized to obtain the sequencing libraries and 4.) triplicate PCR reactions with six cycles of PCR were used for library enrichment. The PCR reactions were purified using Ampure XP beads at a ratio of 0.9X. The final clean libraries were quantified using Qubit, followed by agarose gel electrophoresis to confirm the fragment size. The sequencing was performed on a single lane of the Illumina NovaSeq 6000 using the SP flowcell with paired-end chemistry 2 × 150bp.

### Genome assembly

Quality of the sequencing data was assessed by FastQC v0.11.9 (Andrews, 2010), and cutadapt v4.6 (Martin, 2011) was used to remove any adapters or low-quality sequences. Prior to assembly, k-mer profiling was done using Meryl v1.3 (https://github.com/marbl/meryl) and GenomeScope v2.0 (Vurture et al., 2017; Ranallo-Benavidez et al., 2020), and assembly parameters were adjusted based on the expected genome size and coverage.

HiFiasm version 0.16.1 (Cheng et al., 2021) was used to assemble the PacBio HiFi reads using the arguments -m 10000000 -p 100000 –hom-cov 20, and –hic1 –hic2 to integrate the Hi-C read data and produce two raw assemblies. Statistics and k-mer profiles of both assemblies were checked using gfastats v1.3.6 (Formenti et al., 2022) and Merqury v1.3 (Rhie et al., 2020), and the most complete assembly was chosen as primary, with the other kept as alternate. Duplicate contigs were removed using purge_dups v1.2.6 (Guan et al., 2020) from both assemblies. We then continued with the scaffolding of the assemblies. To process the HiC data, we followed the base pipeline by ArimaGenomics (https://github.com/ArimaGenomics/mapping_pipeline). Briefly, we mapped the HiC data to the genome assemblies using bwa-mem2 v2.2.1 (Vasimuddin et al., 2019). The mapped reads were then parsed and filtered using samtools v1.18 (Danecek et al., 2021) and the scripts provided on the ArimaGenomics github page above, and duplicate reads were removed using the MarkDuplicates tool of the GATK toolkit v4.3.0.0 (O’Connor & van der Auwera, 2020).

For scaffolding, YaHS v1.2 (Zhou et al., 2023) was run using parameters -e GATC -l 5000 –no-contig-ec for the assembly and the filtered Hi-C bam file. Contiguity and general genome statistics were calculated using QUAST v5.2.0 (Mikheenko et al., 2018). We assessed the completeness of the genome by calculating the number of complete single copy orthologs with BUSCO v5.5.0 (Manni et al., 2021), using using both the glires_odb10 and mammalia_odb10 databases.

### Telomere and repeat annotation

Repeat annotation of the genome was performed with RepeatModeler v2.0.5 (Flynn et al., 2020). Using the repeat library produced by RepeatModeler, we masked the scaffolded genome using RepeatMasker v4.1.6 (Smit et al., 2013-2015).

Telomeric sequences were identified using the Telomere Identification ToolKit (tidk v0.2.0) by first running tidk explore to identify potential telomeric repeat sequences, and then tidk search with the sequence of “AACCCT”.

### Manual curation

The assembled and annotated genome was manually curated to further improve its quality as described in (Howe et al., 2021) using PretextView https://github.com/sanger-tol/PretextView/ and the rapid curation workflow https://gitlab.com/wtsi-grit/rapid-curation. The manual curation allows the identification and fixing of erroneous scaffold assemblies and contig duplications.

### Comparison with previous assembly

We performed a comparison of our scaffolded assembly to the current *L. timidus* genome assembly (NCBI Accession number: GCA_009760805) as well as the *L. europaeus* reference genome assembly (NCBI accession number: GCF_033115175.1). Mapping to the genome was performed using minimap2 version 2.21 (Li, 2018) with the arguments *-asm5*. A dot plot of the alignment was created using the R script pafCoordsDotPlotly.R (https://github.com/tpoorten/dotPlotly).

## Results

### Genome assembly

The expected haploid genome size of *L. timidus* is 3.1785 Gbp (Vinogradov, 1998), containing 23 autosomal chromosomes as well as X and Y sex chromosomes (Gustavsson, 1972). PacBio HiFi sequencing produced reads with sequence N50 – the length of the shortest read at 50% of the total sequence length - of 19.97 kb with 21× coverage. Based on the PacBio HiFi data, the expected genome size using k-mer length k =28 is 2.65 Gbp (Figure 2A). The data enabled haplotype-phasing of a primary and an alternative assembly (Figure 2B). The illumina sequencing of the Hi-C data produced 995,011,507 paired reads representing about 56 × coverage of the genome. The duplication rate of the Hi-C data was 18 %. Assembly with HiFiasm yielded a primary contig assembly of 2,838,807,612 bp made up of 2332 contigs and an alternate assembly of 2,654,293,704 bp of 1989 contigs (The longest contigs for the assemblies were 33 Mbp and 26.6 Mbp, and the contig N50s 4.7 Mbp and 4.8 Mbp, respectively. Using the uniquely mapped Hi-C data, we were able to scaffold the contigs and fix misassembled contigs. The primary Hi-C scaffolded assembly was 2,654,826,856 bp in size, the largest scaffold was 181 Mbp and the scaffold N50 127 Mbp.

**Figure 2.**
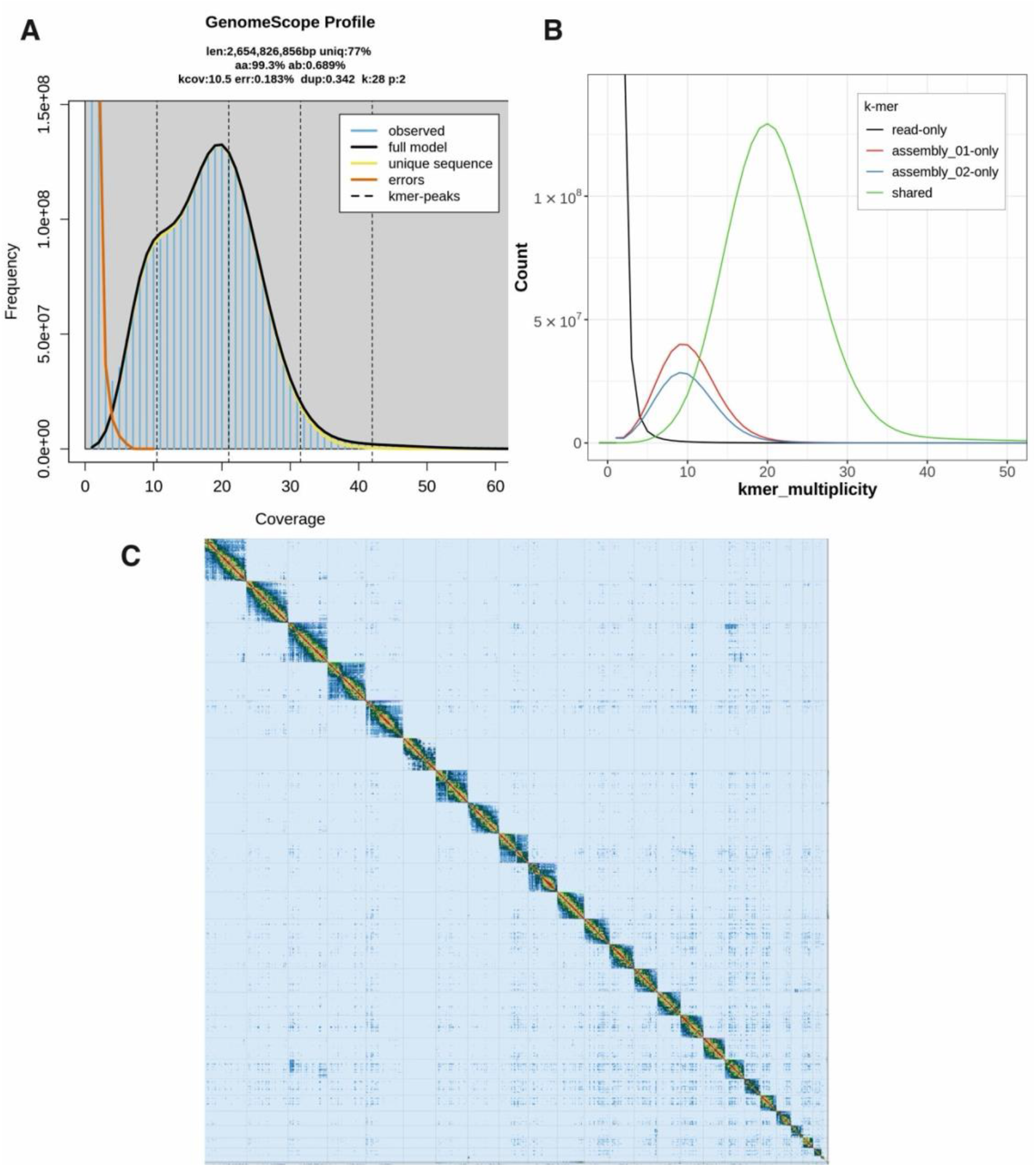
An overview of the mountain hare genome assembly. A) K-mer frequency and coverage distribution obtained from the GenomeScope analysis. The observed k-mers (blue lined area) include any and all k-mers observed. K-mers close to coverage 1 are putative errors (orange outline) due to technical errors in sequencing. Dotted lines show peaks for each ploidy: at the haploid, or heterozygous peak, unique k-mers were found enough times for haploid coverage (10.5), at the diploid, or homozygous peak, k-mers were found enough times for diploid coverage (21). The triploid and tetraploid coverage is also marked, although there are no obvious peaks there, as expected for a known diploid species. K-mers at this location in a diploid assembly represent duplicated heterozygous and duplicated homozygous regions. B) k-mer distribution profile after the first assembly step with Hifiasm showing also the amount of unique k-mers for the primary (01) and alternative (02) assemblies. Shared k-mers are found in both assemblies, while read-only k-mers are sequencing errors, and not found in either assembly. C) Contact map of Hi-C scaffolding of the primary assembly after manual curation, generated with PretextView.

To further improve the assembly, manual curation was performed at the Sanger Institute using Hi-C maps (Figure 2C). The manual curation changed the scaffold N50 of 127.6 Mb to 125.8 Mb and caused a reduction in scaffold count of 103 to 85 in the primary and 128 to 109 in the alternate assembly. Of the finalized assembly, 93.16 % could be assigned to 25 identified chromosomes (23 autosomes plus X and Y) (Table 1). Autosomes were named according to their size. After decontamination and manual curation, the scaffold counts are 85 for the primary, and 109 for the alternate assembly. Detailed final assembly statistics are shown in Table 2. The curated genome length was 2,695,305,354 bp. The mitochondrial genome assembly and the annotation of the functional loci have been presented elsewhere (Tapanainen et al., 2024).

**Table 1.** Sequence assignment of the primary mountain hare genome assembly.

|  | Length | % of total length | Count |
| --- | --- | --- | --- |
| **Autosomes** | 2,519,453,057 | 93.48 | 23 |
| **Unlocalized on autosomes** | 1,279,896 | 0.05 | 1 |
| **Other scaffolds** | 11,202,567 | 0.42 | 58 |
| **X** | 139,721,860 | 5.18 | 1 |
| **Y** | 24,927,870 | 0.92 | 1 |
| **MT** | 17,482 |  | 1 |
| **TOTAL** | 2,695,305,354 |  | 85 |

**Table 2.** Assembly statistics for the Hi-C scaffolded, post-curation sequencing data for the mountain hare reference genome.

| Assembly statistics | Primary | Alternate |
| --- | --- | --- |
| # scaffolds | 85 | 109 |
| Total scaffold length | 2,695,305,354 | 2,556,802,277 |
| Average scaffold length | 31,709,474.75 | 23,456,901.62 |
| Scaffold N50 | 125,755,317 | 126,936,470 |
| Scaffold auN | 125,197,910.08 | 126,023,208.11 |
| Scaffold L50 | 9 | 9 |
| Largest scaffold | 182,608,553 | 185,135,719 |
| Smallest scaffold | 10,771 | 21,369 |
| # contigs | 1,414 | 1,454 |
| Total contig length | 2,695,169,802 | 2,556,665,250 |
| Average contig length | 1,906,060.68 | 1,758,366.75 |
| Contig N50 | 4,885,811 | 5,114,768 |
| Contig auN | 6,632,874.02 | 6,902,786.36 |
| Contig L50 | 157 | 136 |
| Largest contig | 32,947,725 | 26,587,001 |
| Smallest contig | 967 | 21,369 |
| # gaps in scaffolds | 1,329 | 1,345 |
| Total gap length in scaffolds | 135,552 | 137,027 |
| Average gap length in scaffolds | 102.00 | 101.88 |
| Gap N50 in scaffolds | 100 | 100 |
| Gap auN in scaffolds | 104.10 | 103.78 |
| Gap L50 in scaffolds | 650 | 660 |
| Largest gap in scaffolds | 200 | 200 |
| Smallest gap in scaffolds | 26 | 27 |
| Base composition (A:C:G:T) |  |  |
| A | 758,096,831 | 716,813,053 |
| C | 589,442,155 | 561,406,584 |
| G | 589,603,553 | 561,320,925 |
| T | 758,027,263 | 717,124,688 |
| GC content % | 43.75 | 43.91 |
| # soft-masked bases | 0 | 0 |
| # segments | 1,414 | 1,454 |
| Total segment length | 2,695,169,802 | 2,556,665,250 |
| Average segment length | 1,906,060.68 | 1,758,366.75 |
| # gaps | 1,329 | 1,345 |
| # paths | 85 | 109 |

The BUSCO scores of the *L. timidus* assembly suggest that it is near-complete, with the following results: Complete: 95.1 % [Single copy: 92.3 %, Duplicated: 2.7 %] Fragmented 0.8 %, and Missing 4.1 % based on the mammalia_odb10 database of mammalian orthologs and Complete: 93.2% [Single copy: 90.6 %, Duplicated: 2.7 %], Fragmented 1.1 % and Missing 5.6 % based on the glires_odb10 database of lagomorph orthologs. The categories refer to if a gene’s ortholog, that was expected to be found in a single copy, was found completely, partially (fragmented), in a single copy or in multiple copies, or if it was completely missing. The number of total groups searched were 9226 and 13798, respectively (Table 3, Figure 3). T-antigen vector insertions were detected on the chromosomes 17 and 19. The insertions were in intergenic regions and have been replaced with equal length of Ns in the final assembly. The SV40 T-antigen-containing vector is a recombinant expression construct used to immortalize the fibroblast cell lines (Gaertner et al., 2023). Immortalization simplifies handling for subsequent experiments and minimizes phenotypic variation between cell lines but is not required for genomic work.

**Table 3.** BUSCO completeness statistics for the mountain hare reference genome assembly.

| **BUSCO** | **Primary** |  | **Alternate** |  |
| --- | --- | --- | --- | --- |
| **Mammalia** | **Count** | **%** | **Count** | **%** |
| Complete BUSCOs (C) | 8770 | 95.06 | 8601 | 93.23 |
| Complete and single-copy BUSCOs (S) | 8517 | 92.32 | 8378 | 90.81 |
| Complete and duplicated BUSCOs (D) | 253 | 2.74 | 223 | 2.42 |
| Fragmented BUSCOs (F) | 76 | 0.82 | 84 | 0.91 |
| Missing BUSCOs (M) | 380 | 4.12 | 541 | 5.86 |
| Total BUSCO groups searched | 9226 |  | 9226 |  |
| **Glires** |  |  |  |  |
| Complete BUSCOs (C) | 12863 | 93.22 | 12553 | 90.98 |
| Complete and single-copy BUSCOs (S) | 12495 | 90.56 | 12226 | 88.61 |
| Complete and duplicated BUSCOs (D) | 368 | 2.67 | 327 | 2.37 |
| Fragmented BUSCOs (F) | 157 | 1.14 | 149 | 1.08 |
| Missing BUSCOs (M) | 778 | 5.64 | 1096 | 7.94 |
| Total BUSCO groups searched | 13798 |  | 13798 |  |

**Figure 3.**
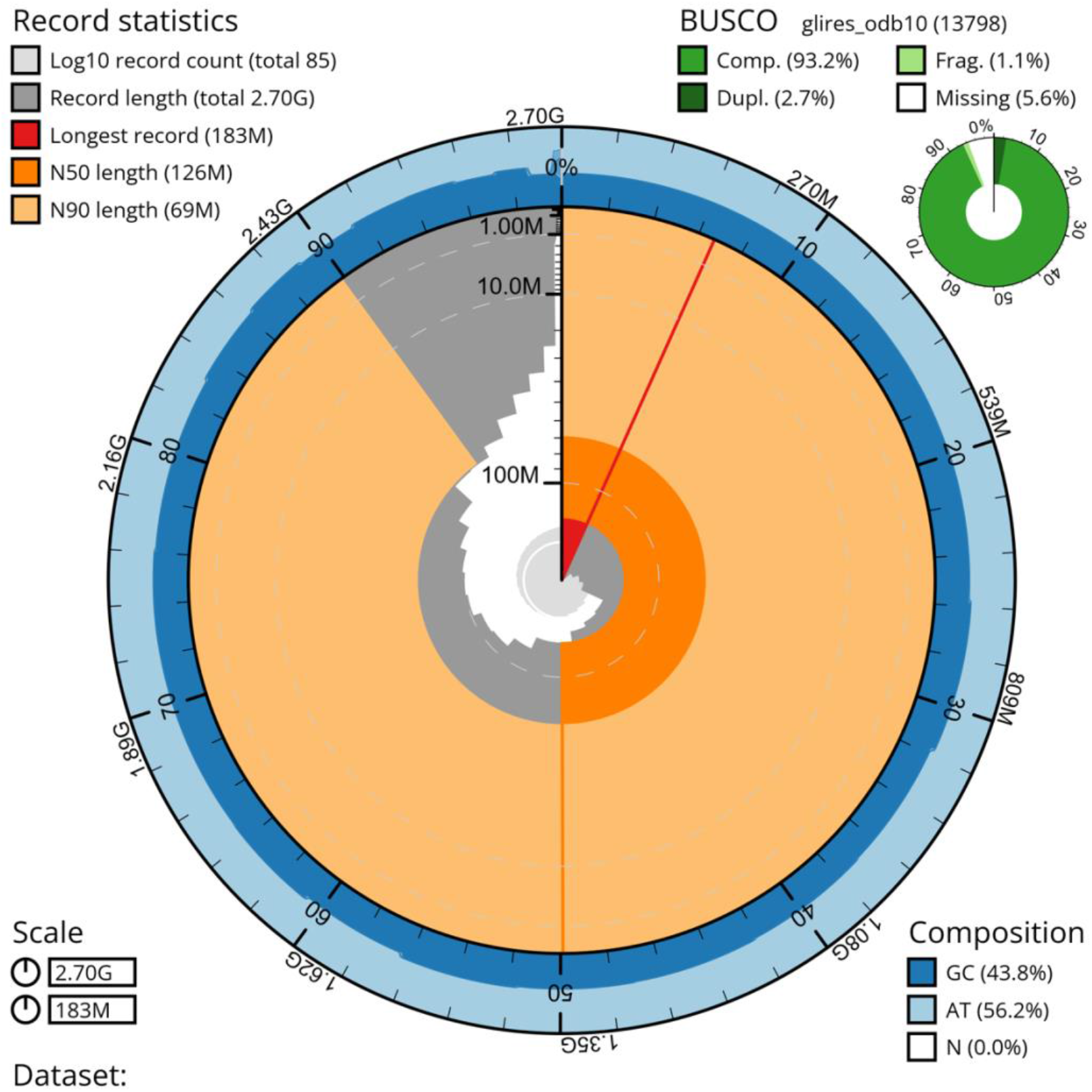
A snail plot summary of assembly statistics for the final, manually curated 2,695,305,354 bp primary assembly. The circumference of the plot symbolizes the full genome length of 2.7 Gbp. The main plot is divided into 1,000 size-ordered bins around the circumference with each bin representing 0.1 % of the final assembly. The distribution of sequence lengths is shown in dark grey with the plot radius scaled to the longest sequence present in the assembly (182,608,553 bp, shown in red). Due to the data being divided into bins, the sequence lengths seem to change in a jagged pattern instead of a smooth one. Orange and pale-orange arcs show the N50 and N90 sequence lengths (125,755,317 and 68,953,458 bp), respectively, while the red arc signifies the length of only one record, the longest one. The scale starts at the outer side, and it is logarithmic. The pale grey spiral shows the cumulative sequence count on a log scale with white scale lines showing successive orders of magnitude. The blue and pale-blue area around the outside of the plot shows the distribution of GC, AT and N percentages in the same bins as the inner plot.A summary of complete, fragmented, duplicated and missing BUSCO genes based on the glires_odb10 set is shown in the top right.

Repeat detection with RepeatModeler produced a curated custom repeat library of 28,386 unique repetitive elements classified into 809 repeat families. Interestingly, the proportion of the genome masked as repetitive elements was very high, at 42.35 % of the genome, similar to the situation observed with our brown hare genome assembly (46.8 %) (Michell et al., 2024). In contrast, in the Irish hare genome assembly (GCA_009760805), only 28 % of the genome was masked (Marques et al., 2020). The *k*-mer based estimate of the repetitive elements in our mountain hare was 23.0 %, which is slightly less than the 26.7 % observed for the brown hare (Michell et al., 2024).

Telomeric sequences were found at both ends of 9 chromosomes, and only on one end of 10 more chromosomes (Figure 4). More than half of the chromosomes (13) also contain telomeric elements not only at their ends but also at various internal locations.

**Figure 4.**
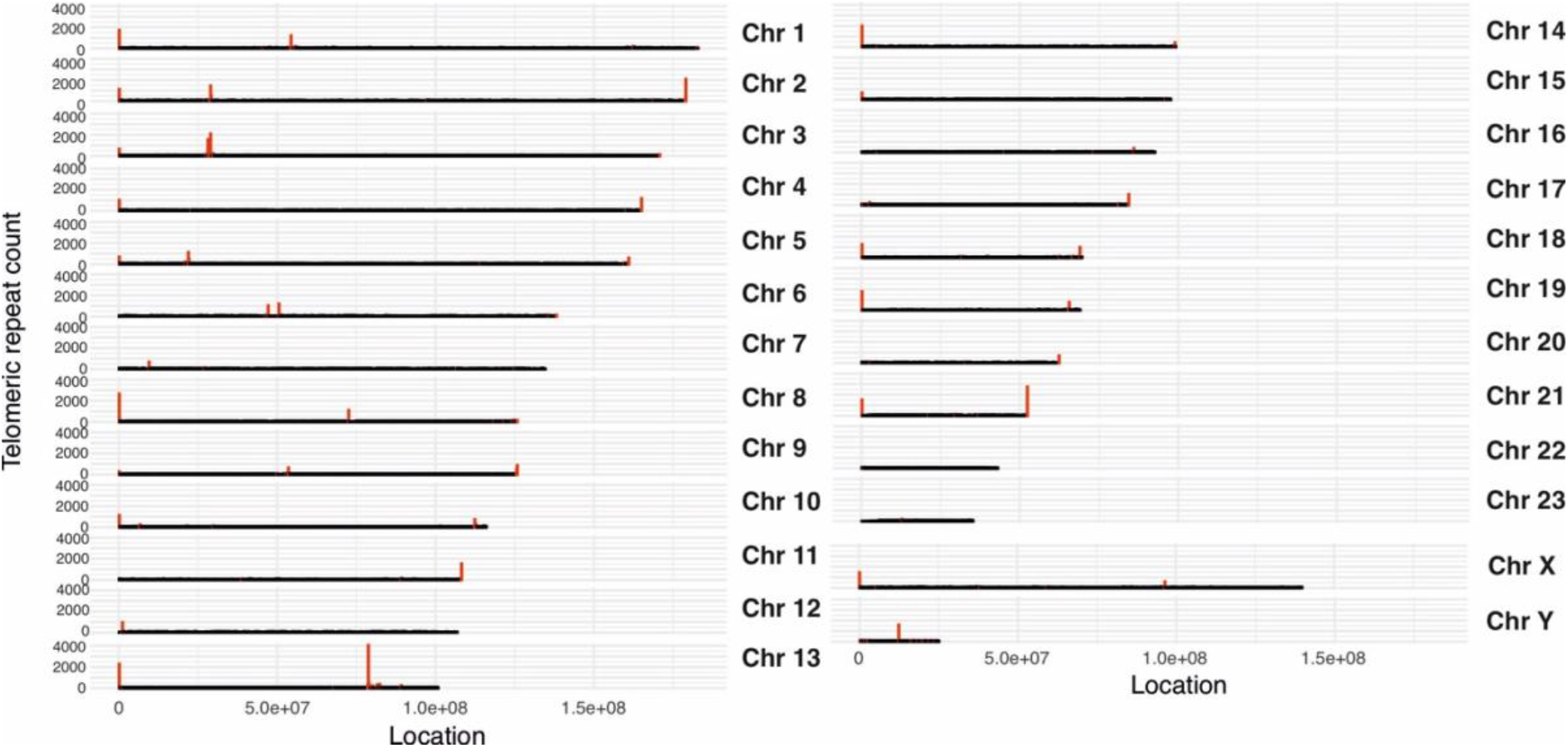
Telomeric repeat arrays on the mountain hare chromosome assemblies, calculated in windows of 200 kb. The location and relative length of the repeat is presented by the red bars. Short telomeric repeats are not visible in the graph.

The genome assemblies can be accessed via BioSample accession SAMN41430840 as well as BioProject accession PRJNA1112569 for the primary assembly and raw data of the PacBio and Hi-C sequencing, and PRJNA1112568 for the alternative assembly.

### Comparison to previous assemblies

Minimap2 was able to align 99.95 % (97.57 % primary mapped) of the contig sequences from the Irish hare genome assembly with the presented mountain hare genome assembly (Figure 5A). The two assemblies have a high level of sequence similarity as well as a high degree of synteny. As pointed out earlier, the Irish hare genome assembly retains the chromosomal arrangement of the rabbit genome (Beklemisheva et al., 2011) used for its scaffolding (Marques et al., 2020). As a result, a few notable differences are present in the chromosome assignments, which were also noted when comparing the Irish hare assembly with the brown hare assembly (Michell et al., 2024). Specifically, hare chromosomes Chr7 and Chr12 correspond to Chr1 of the rabbit genome, and chromosomes Chr13 and Chr16 correspond to rabbit Chr2 (Figure 5A), explaining the karyotype difference (2n = 44 vs. 2n = 48).

**Figure 5.**
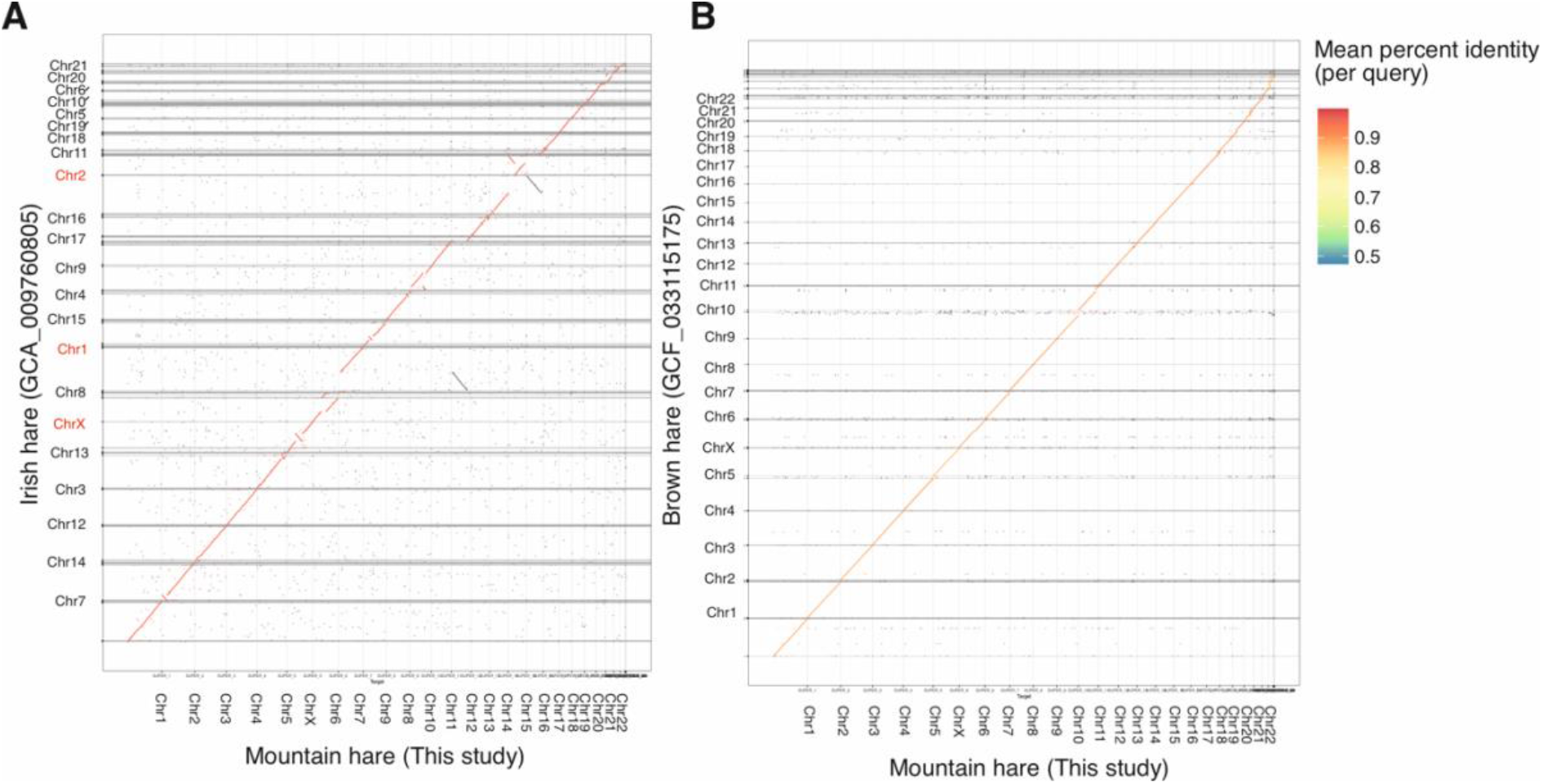
A comparison of the LT1 mountain hare (*Lepus timidus timidus*) genome assembly with A) the previous Irish hare (*Lepus timidus hibernicus*) pseudoreference assembly (GCA_009760805, examples of karyotype differences are highlighted in red) and B) our previously published European brown hare (*Lepus europaeus*) genome assembly (GCF_033115175). Note the higher synteny but lower sequence similarity compared to (A). See the Results section for further explanation.

Consequently, the brown hare genome (Michell et al. 2024, NCBI RefSeq GCF_033115175.1) shows an even higher degree of synteny with our mountain hare genome assembly (Figure 5B), with Minimap2 aligning 99.87 % (78.77 % primary mapped) of the sequences. However, while the visual representation of the two genomes does not show major rearrangements, the relatively lower percentage of primary mapped contigs with minimap2 also indicates a higher occurrence of chimeric alignments, suggesting minor differences in chromosome sequence organization. Despite the high levels of synteny, the sequence similarity between the brown hare and mountain hare assemblies is lower than that between the mountain hare subspecies. This is illustrated by the sequence identity colors in the alignments (Figure 5A vs. B). The mean sequence similarity, as calculated from the number of matches between each of the aligned scaffolds, is 96.77 % for the mountain hare vs. Irish hare and 93.09 % for the mountain hare vs. brown hare.

## Discussion

To complement our previously published brown hare reference genome assembly (Michell et al., 2024), we have sequenced and produced a reference-quality genome assembly for the mountain hare. Although large and complex mammalian genomes can pose problems for assembly, our final genome assembly is highly continuous (85 scaffolds in the primary assembly, with an N50 of 126 Mbp, and contig N50 4.9 Mbp) and complete (BUSCO complete 95.1 % mammalia_odb10) (Tables 1–3, Figures 1–3). The genome assembly is at chromosome scale, with all expected 23 autosomes as well as X and Y scaffolded. Our assembly meets the 6.C.Q40 quality criteria for a reference-standard genome assembly (>1 Mb N50 contig continuity and chromosomal scale scaffolding), as proposed by the Earth BioGenome Project (Lawniczak et al., 2022). With the 99.9 % accuracy of PacBio HiFi reads (Wenger et al., 2019) and 21× genome coverage, the assembly also satisfies the reference genome standard of a nucleotide error rate of no more than 1/10,000 (Lawniczak et al., 2022). Most of the chromosome assemblies represent telomere-to-telomere continuity (Figure 4). High copy number telomeric sequences were also present within interstitial positions of some chromosomes, confirming previous observations using FISH staining of mountain hare cells (Forsyth et al., 2005). These regions might be remnants of ancestral chromosome translocation events that gave rise to the current 2n = 46 karyotype in hares.

At approximately 2.7 Gbp the curated genome length is notably shorter than the 3.1785 Gbp estimated for the haploid genome size of *Lepus timidus* from flow cytometry data (Vinogradov, 1998). However, the size of our genome assembly is very similar to the previous pseudoreference assembly (also 2.7 Gbp). We consider it likely that genome assemblies derived from long-molecule sequencing data provide more accurate genome size estimates than older methods such as flow cytometry, which might be influenced by the choice of references and chromosomal organization of the analyzed species. As in the case of our brown hare genome assembly (Michell et al., 2024), we utilized living cultured fibroblasts to overcome the technical challenges in obtaining large quantities of intact, high molecular weight DNA for HiFi sequencing, essential for high quality and highly continuous genome assemblies.

Importantly, our mountain hare genome assembly did not show any genomic rearrangements or aneuploidies that would have been obtained during cell culture. Although genomic changes can occur during cell culture (Didion et al., 2014), these are more likely in cancer cell lines with compromised DNA repair and cell proliferation pathways (He et al., 2023). Genetic changes need time as they appear by chance and increase their prevalence through drift. Therefore, long-term culturing of the cells should be avoided, and low-passage number cells should be preferred for analyses. Ultimately, the genomic stability of any cell line can be only assessed through karyotyping. Here too, modern sequencing approaches offer far greater insight than traditional chromosome-spreading methods. In this context, our hare genome assemblies can serve as benchmarks for using fibroblast cell lines for such work.

However, despite being devoid of karyotypic changes, we noted the insertion of the T-antigen vector used for immortalization of the cells (Gaertner et al., 2023) on chromosome 17 and 19. Interestingly, such insertions are not present in our brown hare genome assembly (Michell et al., 2024), indicating that the immortalization of this brown hare cell line has not worked. Although the insertions were in the intergenic region and we have removed the vector sequences from the final genome assembly, we recognize that this is not an ideal situation as we cannot control for possible insertional effects on the surrounding chromosomal region. Therefore, we recommend that primary cell lines should be used for reference genome work. For other types of studies it needs to be taken into consideration that non-immortalized cell lines have limited lifetime and can show signs of senescence, which can influence analyses.

The presented mountain hare genome assembly shares a high degree of similarity with the existing Irish hare pseudo-reference genome assembly. The observed differences in synteny (Figure 5A) can be attributed to use of the karyotypically different European rabbit (*Oryctolagus cuniculus)* for the chromosomal scaffolding of the Irish hare genome assembly (Marques et al., 2020). Compared to the recently published brown hare genome assembly (Michell et al., 2024), the presented mountain hare genome assembly shows almost identical synteny (Figure 5B), reflecting their close evolutionary relationship. However, when compared to the Irish hare, mountain hare – brown hare comparison yielded a higher number of chimeric alignments, likely due to minor differences in chromosome structure. These structural differences are unlikely to have a major biological significance, given that the two species can hybridize and produce fertile offspring.

The 93.09 % sequence similarity of the mountain hare and brown hare genomes is interesting to contrast with the 96.77 % similarity between the mountain hare subspecies. It should be noted that these similarities are based on matches between scaffold alignments, and comparing specific elements, such as coding sequences alone, would likely yield higher similarity percentages. Nonetheless, these figures highlight the relative differences between species and subspecies genomes, particularly as the Irish hare is known to be genetically distinct from Fennoscandian mountain hares (Giska et al., 2022). More detailed population-level comparisons might allow to pinpoint functional loci contributing to local adaptations (Giska et al., 2019) as well as shed light onto speciation processes (Gaertner et al., 2023).

The mountain hare is an iconic boreoalpine mammalian species with unique physiological and morphological adaptations to cold, snowy winter conditions, it is also an important part of the local food webs. Consequently, the population genetics, evolution and genomics of mountain hares has been intensively studied [e.g. (Melo-Ferreira et al., 2009; Smith et al., 2017; Levänen, Thulin, et al., 2018; Marques et al., 2020; Ferreira et al., 2021; Pohjoismäki et al., 2021; Giska et al., 2022; Michell et al., 2022; Gaertner et al., 2023)]. Together, this genome and the previously published brown hare genome assembly will provide a solid base for future studies on hares. Annotated genome assemblies also enable effective use of reverse genetics, allowing experimental manipulation of any gene or functional feature to study its phenotypic effects. This capability opens opportunities to experimentally demonstrate the functional significance of genetic variants controlling interesting traits, which otherwise could only be indirectly inferred from population genetic analyses. For population genetics, chromosomally scaffolded genome assemblies enable fast and reliable identification of informative SNPs and their linkage. Specifically, the mountain hare genome assembly may offer crucial insights for the conservation of threatened subspecies such as the heath hare (*Lepus timidus sylvaticus*) (C. G. Thulin et al., 2021; Michell et al., 2022) and Irish hare (*Lepus timidus hibernicus*) (Reid, 2011; Caravaggi et al., 2017).

In conclusion, the successful sequencing and assembly of the mountain hare genome, alongside the previously published brown hare genome, mark significant advancements in our understanding of these species’ genomic landscapes. These high-quality genome assemblies not only provide invaluable resources for exploring the genetic basis of adaptation and resilience in hares, but also pave the way for future research in evolutionary biology, conservation genetics, and ecological genomics.

## Acknowledgements

We acknowledge the DNA Sequencing and Genomics Laboratory, Institute of Biotechnology, University of Helsinki for the PacBio and Hi-C sequencing. We are grateful to the European Reference Genome Atlas (ERGA) for the opportunity to include our mountain hare genome into their pilot project. Mr Jussi Mähönen, MSc Manu Soininmäki and Finnish hound Hupi are thanked for their contribution in obtaining the mountain hare specimen. We are indebted also to Ms Anita Kervinen for her valuable laboratory assistance and Dr. Mervi Kunnasranta for kindly allowing to use her mountain hare photo in the publication.

## Data, scripts, code, and supplementary information availability

The genome assembly and sequencing data is available from the NCBI database under BioProject ID PRJNA1112569 and PRJNA1112568 and accession numbers JBEDNU000000000 and JBEDNT000000000. The cell line is available through the corresponding author as well as from the ZFMK biobank.

